# Longitudinal Increases in Ventral Attention Network Connectivity Predict Heightened Reward and Punishment Sensitivities via Self-Control Decline in Emerging Adulthood

**DOI:** 10.64898/2026.11.25.769126

**Authors:** Jingzhen He, Haichao Zhao, Xu Lei, Jiang Qiu, Tingyong Feng, Hong Chen, Ofir Turel, Qinghua He

**Author notes:** Corresponding author at: Faculty of Psychology, Southwest University, 2 Tiansheng Rd, Chongqing 400715, China. E-mail address (Q. He).

## Abstract

Emerging adulthood is a critical developmental window for the maturation of executive control and the reorganization of intrinsic brain networks. However, the longitudinal neurobiological mechanisms underlying changes in cognitive control—and their associations with trait sensitivities—remain poorly understood. In a two-year longitudinal study, 332 emerging adults underwent behavioral assessments and resting-state functional magnetic resonance imaging (fMRI). Latent Change Score Modeling (LCSM) revealed that longitudinal increases in reward and punishment sensitivities were primarily predicted by the progressive decline in self-control, rather than baseline or concurrent impulsivity. Neurally, by targeting the macroscopic connectome utilizing rigorous change score models adjusted for baseline variances and longitudinal head motion, we found that longitudinal increases in intra-network connectivity within the Ventral Attention Network (VAN) specifically and robustly tracked this self-control decline. Furthermore, a unified path analysis revealed a robust full mediation mechanism: heightened VAN intra-network connectivity predicted increased environmental sensitivities strictly through the decline in self-control. These findings identify increased VAN connectivity as a specific neural marker of diminished cognitive control, highlighting the preservation of self-control capacity as a key mechanistic target for promoting adaptive resilience during emerging adulthood.

## 1. Introduction

Impulsivity and reward and punishment sensitivity are individual differences that relate to various mental health challenges, including addiction, anxiety, and depression (Carver & White, 1994; Maxwell, Gardiner, & Loxton, 2020; Moeller, Barratt, Dougherty, Schmitz, & Swann, 2001). *Reward and punishment sensitivity* refers to an individual’s ability to respond to reward and punishment stimuli (Torrubia, Avila, Moltó, & Caseras, 2001). According to Reinforcement Sensitivity Theory (RST), an individual’s sensitivity to rewards and punishments is regulated by the Behavioral Activation System (BAS) and the Behavioral Inhibition System (BIS), respectively (Gray, 1970). These systems influence how individuals respond to environmental stimuli. *Impulsivity* is another individual difference, capturing an individual’s lack of careful consideration during decision-making as manifested in a tendency for quick reactions (Patton, Stanford, & Barratt, 1995). As a significant manifestation of BAS activation, impulsivity can result in individuals being more prone to engage in risky and poorly thought-out behaviors, driven by immediate rewards (Quilty & Oakman, 2004).

Self-control is a cognitive regulatory capability, capturing the ability to regulate impulses and control behavior in order to achieve long-term goals (J. P. Tangney, Baumeister, & Boone, 2004). When self-control is used continuously or repeatedly, this resource may become temporarily reduced (i.e., people experience ego depletion), thereby impairing subsequent self-regulation (Baumeister, Vohs, & Tice, 2007). Notably, diminished self-control can be associated with increased impulsive behaviors (De Ridder, Lensvelt-Mulders, Finkenauer, Stok, & Baumeister, 2012; Heatherton & Tice, 1994; Lin & Feng, 2024; Sharma, Markon, & Clark, 2014). At the same time, there is evidence that individuals with higher self-control also exhibit lower sensitivity to rewards. This suggests that greater self-control can effectively reduce the pursuit of immediate gratification, facilitate the perseverance toward long-term goals, and promote more rational decision-making (Duckworth & Steinberg, 2015; Mischel, Shoda, & Rodriguez, 1989). While the original strength model conceptualizes ego-depletion as a temporary state following short-term exertion, less is known about whether sustained cognitive demands over developmental periods (such as emerging adulthood) relate to long-term trait changes. Despite the importance of these factors for healthy decision-making, their longitudinal interactions remain poorly understood. Specifically, we aim to explore whether long-term reductions in self-control capacity might similarly associate with altered environmental sensitivities over time, beyond the effects of baseline impulsivity.

To elucidate the physiological anchors of this cognitive change, we turn to network neuroscience, which offers valuable insights into how the macroscopic functional organization of the brain supports complex behavior (Bullmore & Sporns, 2009; Stam & Reijneveld, 2007; Van Den Heuvel & Pol, 2010). Human cognition and self-regulation are governed by complex interactions within large-scale functional networks, rather than isolated regions (Friston, 2011). To systematically investigate these systems without a priori spatial bias, the present study employs the widely validated seven-network parcellation (Yeo et al., 2011). Accordingly, the functional architecture of the brain is delineated into the Visual Network (VN), Somatomotor Network (SMN), Dorsal Attention Network (DAN), Ventral Attention Network (VAN), Limbic Network (LN), Frontoparietal Network (FPN), and Default Mode Network (DMN). To operationalize the functional integrity of these macroscopic systems, we quantify their internal communication through resting-state functional connectivity (FC). Specifically, the strength of FC within these canonical networks—often termed intra-network integration—reflects the intrinsic neural coordination essential for maintaining cognitive functioning and emotional regulation (Finn & Bandettini, 2021; van den Heuvel & Sporns, 2019).

While previous studies have primarily linked static, cross-sectional network architectures to psychiatric vulnerabilities, the longitudinal topological reconfiguration of these networks remains underexplored (Preti, Bolton, & Van De Ville, 2017). Importantly, the human brain functions as a competitively interacting system constrained by limited metabolic and computational resources (Bullmore & Sporns, 2012). Consequently, longitudinal shifts in intra-network integration—such as a specific subnetwork exhibiting increased connectivity over time—may relate to changes in cognitive allocation. Such increased intra-network connectivity may relate to a redistribution of cognitive resources, potentially constraining top-down executive systems (Menon, 2011).

Despite these theoretical links, empirical investigations have largely relied on cross-sectional designs, leaving the longitudinal neurobehavioral associations unresolved. To address this critical gap, the current study integrates Latent Change Score Modeling (LCSM) with two-wave longitudinal resting-state fMRI. We propose a unified longitudinal competitive framework guided by two primary hypotheses.

Behaviorally, we hypothesize that longitudinal reductions in self-control will predict concurrent increases in both reward and punishment sensitivities over time. Neurally, without constraining our focus a priori, we systematically track the topological reconfiguration across all seven canonical subnetworks. We hypothesize that increased intra-network connectivity within networks governing bottom-up attentional capture will robustly couple with this regulatory decline. Ultimately, we aim to validate a full longitudinal mediation pathway wherein longitudinal neural reconfiguration alters environmental sensitivities via the decline in self-control capacity. An overview of the multi-modal longitudinal study design and the unified analytical framework is illustrated in Figure 1.

**Figure 1.**
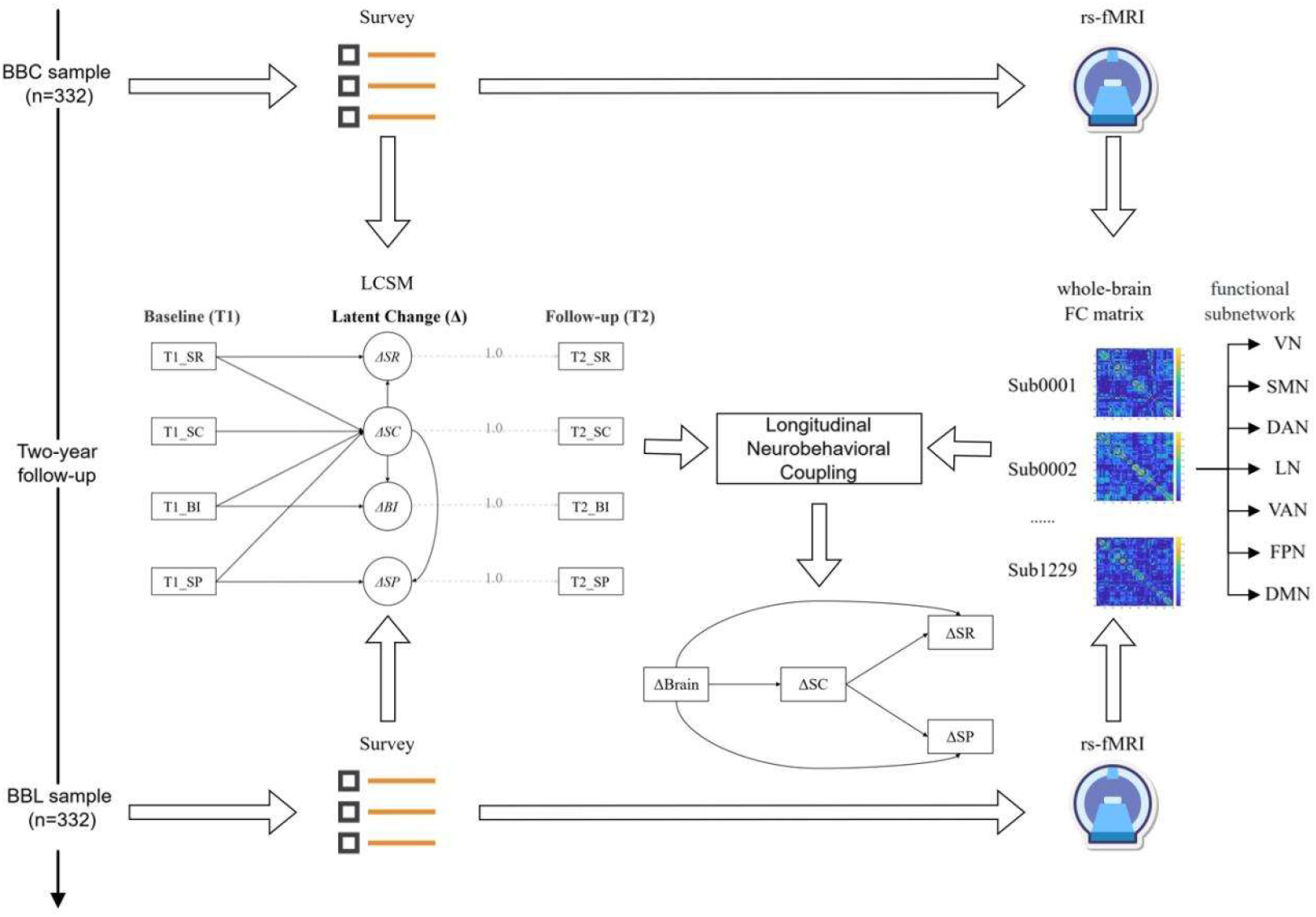
Global analytical framework and multi-modal longitudinal study design. The schematic delineates the comprehensive pipeline integrating behavioral and neuroimaging data across a two-year longitudinal follow-up. Δ represents the raw longitudinal change scores from Time 1 to Time 2. SR = Sensitivity to Reward; SP = Sensitivity to Punishment; SC = Self-Control; BI = Barratt Impulsiveness; VN = Visual Network; SMN = Somatomotor Network; DAN = Dorsal Attention Network; LN = Limbic Network; VAN = Ventral Attention Network; FPN = Frontoparietal Network; DMN = Default Mode Network; LCSM = Latent Change Score Modeling; FC = Functional Connectivity.

## 2. Methods and measures

### 2.1. Participants

To investigate the longitudinal neurobehavioral associations, the current study employed a two-wave longitudinal design spanning approximately two years. Initial recruitment and baseline data collection (Time 1) occurred between September 2019 and December 2019, during which university students were recruited via online platforms, email invitations, and campus advertisements to complete behavioral assessments and an 8-minute resting-state fMRI (rs-fMRI) scan. A comprehensive follow-up re-assessment (Time 2) was subsequently conducted from September to December 2021.

To construct a rigorous longitudinal neurobehavioral dataset, strict multi-modal inclusion criteria were applied. Participants were required to possess complete, high-quality data across both modalities (behavioral surveys and rs-fMRI) at both Time 1 and Time 2. Individuals with incomplete questionnaire responses or poor-quality MRI data at either time point (e.g., mean framewise displacement exceeding 0.2 mm or more than 30% of timepoints scrubbed) were excluded. Following these stringent quality control measures, a final cohort of 332 individuals was retained for all cross-sectional and longitudinal neurobehavioral coupling analyses. All procedures complied with the Declaration of Helsinki and were approved by the Ethics Committee of Chongqing Ninth People’s Hospital (Approval: 2019 lunshen 016). Participants’ capacity to consent was affirmed based on their legal age of majority and demonstrated comprehension of the study. After a full explanation of the protocol, all participants provided written informed consent. To protect human subjects, data were strictly anonymized, and participants retained the right to withdraw at any time without penalty.

### 2.2. Behavior measures

#### 2.2.1. Sensitivity to Punishment and Sensitivity to Reward Questionnaire (SPSRQ)

The SPSRQ was used to evaluate individuals’ responsiveness to rewarding and punishing stimuli (Torrubia et al., 2001). This instrument includes two main scales: Reward Sensitivity (SR), which reflects the degree of motivation and tendency to pursue rewards, and Punishment Sensitivity (SP), which captures anxiety-related and avoidance reactions triggered by punishment signals. The questionnaire consists of 48 items requiring “Yes” or “No” responses, with even-numbered items assessing SR and odd-numbered items evaluating SP. Each subscale contains 24 items, and the score for each ranges from 0 to 24. In the current sample, the internal consistency (Cronbach’s alpha) at baseline was 0.68 for SR and 0.81 for SP. Furthermore, the longitudinal test-retest reliabilities between the two waves were robust, yielding r = 0.61 (*p* < 0.001) for SR and r = 0.71 (*p* < 0.001) for SP.

#### 2.2.2. Self-Control Scale (SCS)

The SCS captures individuals’ capacity for self-regulation and impulse control in various behavioral and psychological contexts (J. P. Tangney et al., 2004). Here, we used its adaptation to the Chinese cultural context (Tan & Guo, 2008). The revised version includes 19 items measuring self-control, each rated on a 5-point Likert scale ranging from 1 (completely untrue) to 5 (completely true). Total scores range from 19 to 95, with higher scores indicating stronger self-control. The internal consistency (Cronbach’s alpha) for this scale at baseline was 0.75 in the current sample, and the longitudinal test-retest reliability was r = 0.68 (*p* < 0.001).

#### 2.2.3. Barratt Impulsiveness Scale (BIS)

The BIS is a widely used self-report questionnaire designed to measure individual impulsivity (Patton et al., 1995). This scale consists of 30 items rated on a 4-point Likert scale and assesses three related yet distinct subdimensions of impulsivity: Attentional Impulsiveness, Motor Impulsiveness, and Non-planning Impulsiveness.

These subscales respectively reflect impulsive traits in rapid cognitive decision-making, tendency toward unpremeditated actions, and lack of future planning. Each subscale contains 10 items, with raw scores ranging from 10 to 50. Scores are converted to standardized scores ranging from 0 to 100 using the formula: ((sum of raw scores − 10) ÷ 40) × 100. The overall BIS score is calculated as the average of the three standardized subscale scores, with higher scores indicating greater impulsivity. In the current sample, the internal consistency (Cronbach’s alpha) of the BIS at baseline was 0.89, with a robust longitudinal test-retest reliability of r = 0.64 (p < 0.001).

### 2.3. Image acquisition

In this study, all participants completed an 8-minute resting-state functional magnetic resonance imaging (fMRI) session using a 3T SIEMENS PRISMA scanner (Erlangen, Germany). During the scan, they were instructed to keep their eyes open and to avoid focusing on any particular thoughts. Functional images were acquired using a gradient echo-planar imaging sequence, resulting in a total of 240 volumes. The scanning parameters are as follows: repetition time (TR) = 2000 ms, echo time (TE) = 30 ms, field of view (FOV) = 224 × 224, flip angle (FA) = 90 °, slices = 62, thickness = 2 mm, slice gap = 0.3 mm, voxel size = 2 × 2 × 2 mm^3^. Additionally, high-resolution T1-weighted structural images were acquired using the magnetization-prepared rapid gradient echo (MPRAGE) sequence. The scanning parameters are as follows: TR = 2530 ms, TE = 2.98 ms, FOV = 224 × 256 mm^2^, resolution matrix = 448 × 512, FA = 7°, slices = 192, thickness = 1.0 mm, inversion time = 1100 ms, voxel size = 0.5 × 0.5 × 1 mm^3^.

### 2.4. Data Preprocessing

The preprocessing of fMRI data was performed utilizing the SPM12 and CONN v20.b toolboxes. The procedure involved several key steps: temporal realignment, correction for distortions caused by field inhomogeneities, and adjustment for head motion. After these corrections, spatial normalization was applied to align each participant’s high-resolution anatomical images with their functional scans. The anatomical images were then segmented into different tissue types, including gray matter, white matter, and cerebrospinal fluid. Normalization to the Montreal Neurological Institute (MNI) template space was achieved through the Dartel algorithm, followed by spatial smoothing using a Gaussian kernel with a 6 mm full-width at half-maximum (FWHM). During denoising, head motion effects were accounted for by regressing out parameters based on the Friston 24-parameter model (Friston, Williams, Howard, Frackowiak, & Turner, 1996), which incorporates the six motion parameters, their temporal derivatives, and squared terms. Motion-related artifacts were further addressed by scrubbing time points where framewise displacement exceeded 0.2 mm, with flagged volumes and their neighboring scans included as nuisance regressors (Power, Barnes, Snyder, Schlaggar, & Petersen, 2012). To reduce physiological noise and residual motion artifacts, the aCompCor technique was applied, extracting the first five principal components from white matter and cerebrospinal fluid signals (Muschelli et al., 2014). Finally, the data were subjected to linear detrending and band-pass filtering within the 0.008 to 0.09 Hz frequency range.

### 2.5. Statistical Analysis

All statistical analyses and data modeling were performed in R version 4.5.1. The analytical pipeline consisted of three sequential phases: behavioral trajectory modeling, neurobiological targeted localization, and unified pathway modeling.

Behavioral Trajectory Modeling: To investigate the temporal associations among impulsivity, self-control, and reward/punishment sensitivities, a four-variable Latent Change Score Model (LCSM) was constructed using the lavaan package. To ensure robustness against potential non-normality, models were estimated using Maximum Likelihood with robust standard errors (the MLR estimator). Model fit was evaluated using standard indices: the chi-square to degrees of freedom ratio (χ²/df), Comparative Fit Index (CFI), Tucker-Lewis Index (TLI), Root Mean Square Error of Approximation (RMSEA), and Standardized Root Mean Square Residual (SRMR).

Neuroimaging Data Analysis: For the neuroimaging data, we utilized preprocessed resting-state fMRI data to construct the whole-brain functional connectivity (FC) matrix for each participant. To systematically characterize the longitudinal changes in functional connectivity of major functional modules, the whole brain was parcellated into seven canonical networks according to the Yeo et al (2011). template: Visual Network (VN), Somatomotor Network (SMN), Dorsal Attention Network (DAN), Ventral Attention Network (VAN), Limbic Network (LN), Frontoparietal Network (FPN), and Default Mode Network (DMN). Connectivity matrices were aggregated at the subnetwork level to extract the intrinsic intra-network integration for each of the seven functional networks at both Time 1 and Time 2.

Neurobehavioral Coupling and Path Analysis: To evaluate longitudinal neurobehavioral coupling while rigorously addressing the confounding effects of initial individual differences and regression to the mean, we adopted an Analysis of Covariance (ANCOVA)-style change score framework. Raw longitudinal change scores (Δ) were computed by subtracting Time 1 from Time 2 values for both behavioral metrics and intra-network connectivities. We then regressed the raw change in self-control onto the longitudinal integration changes of the functional networks, systematically adjusting for baseline self-control, corresponding baseline network connectivity, sex, and longitudinal head motion changes. The False Discovery Rate (FDR) method was applied to adjust p-values for multiple comparisons across the comprehensive network scans. Finally, a unified longitudinal path analysis (mediation model) was constructed to test whether the effects of longitudinal neural reconfigurations on behavioral sensitivities were statistically mediated by the decline in self-control.

## 3. Results

### 3.1. Participant Characteristics

The cross-sectional behavior-brain sample consisted of 332 participants (254 females) with a mean (SD) age of 18.85 (1.29) years. In this sample, the mean (SD) scores were 12.44 (3.61) for SR, 14.18 (4.71) for SP, 59.83 (10.24) for SC, and 38.70 (9.65) for BI. The longitudinal behavior-brain sample involved a two-year follow-up, yielding mean (SD) scores of 12.00 (3.98) for SR, 13.89 (5.41) for SP, 57.09 (10.52) for SC, and 39.53 (10.05) for BI. To assess the statistical significance of these changes over time, paired t-tests were conducted. Results revealed a significant longitudinal decline in self-control (mean change = −2.74, t = −6.04, *p* < 0.001) and reward sensitivity (mean change = −0.44, t = −2.39, *p* = 0.017). Changes in punishment sensitivity (mean change = −0.29, t = −1.33, *p* = 0.183) and impulsivity (mean change = +0.83, t = 1.79, *p* = 0.074) did not reach statistical significance.

### 3.2. Behavioral Longitudinal Trajectories: Associations Between Self-Control and Sensitivities

To investigate the temporal associations among impulsivity, self-control, and reward/punishment sensitivities over time, a four-variable Latent Change Score Model (LCSM) was constructed. The model demonstrated excellent fit to the data: χ²/df = 1.923, CFI = 0.996, TLI = 0.980, RMSEA = 0.053, SRMR = 0.030.

Regarding proportional changes, the baseline levels of all behavioral variables significantly and negatively predicted their own latent change scores (all *ps* < 0.001), reflecting a typical regression-to-the-mean effect in longitudinal data.

For the cross-lagged predictions (see Table 1), baseline impulsivity (T1_BI) did not significantly predict the longitudinal decline in self-control (ΔSC: β = −0.072, *p* = 0.288). Interestingly, however, higher baseline self-control exerted a significant protective effect against the increases of impulsivity over time (ΔBI: β = −0.211, *p* = 0.003). Crucially, when examining the predictors of reward and punishment sensitivities, our model first evaluated baseline traits and found that baseline impulsivity (T1_BI) did not significantly predict longitudinal changes in either reward (ΔSR: β = −0.064, *p* = 0.419) or punishment sensitivity (ΔSP: β = −0.051, *p* = 0.524). Subsequently, to test whether concurrent state fluctuations better explain these behavioral shifts, the changes in self-control and impulsivity were entered simultaneously to compete for variance. Results revealed that the longitudinal change in impulsivity also failed to predict longitudinal changes in either reward sensitivity (β = −0.021, *p* = 0.745) or punishment sensitivity (β = −0.067, *p* = 0.363). Instead, the decline in self-control emerged as a robust and independent negative predictor.

**Table 1.** Standardized and Unstandardized Path Coefficients for the Behavioral LCSM.

| Outcome Variable | Predictor | B | SE | z | *p* | β |
|---|---|---|---|---|---|---|
| Autoregressive Paths |  |  |  |  |  |  |
| ΔSC | T1_SC | -0.335 | 0.057 | -5.83 | < 0.001 | -0.416<sup>***</sup> |
| ΔSR | T1_SR | -0.359 | 0.05 | -7.25 | < 0.001 | -0.382<sup>***</sup> |
| ΔSP | T1_SP | -0.233 | 0.041 | -5.76 | < 0.001 | -0.277<sup>***</sup> |
| ΔBI | T1_BI | -0.461 | 0.063 | -7.29 | < 0.001 | -0.528<sup>***</sup> |
| Cross-Lagged Paths |  |  |  |  |  |  |
| ΔSC | T1_BI | -0.061 | 0.058 | -1.06 | 0.288 | -0.072 |
| ΔBI | T1_SC | -0.174 | 0.059 | -2.93 | 0.003 | -0.211<sup>**</sup> |
| ΔSR | T1_BI | -0.023 | 0.028 | -0.81 | 0.419 | -0.064 |
| ΔSR | T1_SC | -0.046 | 0.026 | -1.77 | 0.076 | -0.138 |
| ΔSR | ΔBI | -0.008 | 0.026 | -0.33 | 0.745 | -0.021 |
| ΔSR | ΔSC | -0.073 | 0.028 | -2.62 | 0.009 | -0.177<sup>**</sup> |
| ΔSP | T1_BI | -0.021 | 0.033 | -0.64 | 0.524 | -0.051 |
| ΔSP | T1_SC | -0.069 | 0.031 | -2.25 | 0.024 | -0.178<sup>*</sup> |
| ΔSP | ΔBI | -0.031 | 0.035 | -0.91 | 0.363 | -0.067 |
| ΔSP | ΔSC | -0.167 | 0.034 | -4.97 | < .001 | -0.348<sup>***</sup> |

Note. B = unstandardized path coefficient; SE = standard error; z = critical ratio; β = standardized path coefficient. T1 = baseline measure; Δ = latent change score. BI = Barratt Impulsiveness; SC = Self-Control; SR = Sensitivity to Reward; SP = Sensitivity to Punishment. \* *p* ≤ 0.05. \*\* *p* < 0.01. \*\*\* *p* < 0.001.

Specifically, a greater decline in self-control (a smaller ΔSC) was significantly associated with longitudinal changes in reward sensitivity (β = −0.177, *p* = 0.009) and punishment sensitivity (β = −0.348, *p* < 0.001). These results robustly indicate that the changes in reward/punishment sensitivities is not predetermined by baseline impulsive traits, nor is it merely a byproduct of concurrent impulsivity surges; rather, it is primarily driven by the exhaustion of self-regulatory resources during the follow-up period.

The structural architecture of the four-variable LCSM is visually synthesized in Figure 2. For parsimony, non-significant cross-lagged paths and residual variances are omitted. Complementing the parametric details in Table 1, the path diagram explicates the topological configuration of the model. Specifically, it maps the significant baseline covariances and illustrates the formal specification of the latent change scores via fixed constraints. This macroscopic visualization structurally underscores the competitive pathways, confirming the predominant role of self-control decline over impulsivity in predicting the longitudinal increases in reward and punishment sensitivities.

**Figure 2.**
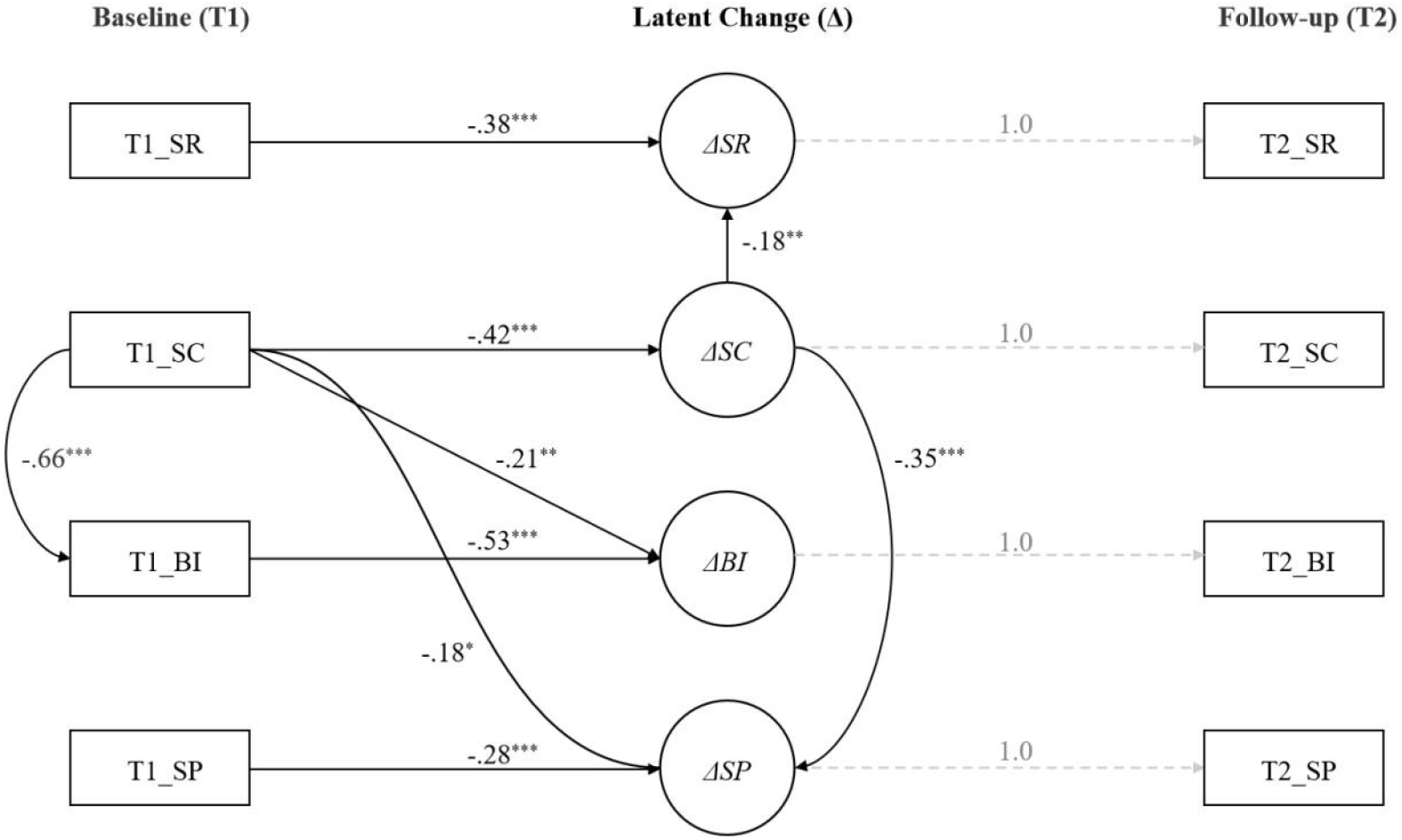
Four-Variable Latent Change Score Model (LCSM) of Self-Control, Impulsivity, and Reward/Punishment Sensitivities. Standardized path coefficients (β) are presented. Rectangles represent observed variables at baseline (T1) and follow-up (T2); circles represent unobserved latent change scores. Solid black lines indicate statistically significant paths. Dashed gray lines represent fixed parameters required for the mathematical identification of the latent change scores. * *p* < 0.05, ** *p* < 0.01, *** *p* < 0.001.

### 3.3. Baseline Prediction and Longitudinal Co-change of Intra-Network Integration in Self-Control Decline

To investigate the role of intra-network integration in longitudinal self-control decline, we first examined neural vulnerability markers at baseline. Results revealed that static baseline functional architecture alone provided limited predictive value. Although specific networks (e.g., Ventral and Dorsal Attention Networks) showed initial predictive trends, no baseline intra-network connectivity robustly predicted subsequent self-control decline after False Discovery Rate (FDR) correction (all *p*_FDR > 0.10; Table 2). This suggests that initial topological architecture is insufficient to predict long-term cognitive resource reduction.

**Table 2.** Baseline Vulnerability and Longitudinal Co-change of Intra-Network Integration in Longitudinal Self-Control Decline.

| Network | Baseline Vulnerability (B) | t | *p*_FDR | Longitudinal Co-change (B) | t | *p*_FDR |
|---|---|---|---|---|---|---|
| Visual Network | 8.07 | 2.2 | 0.106 | 8.18 | 2.24 | 0.358 |
| Somatomotor Network | -1.18 | -0.24 | 0.809 | -2.44 | -0.52 | 0.922 |
| Dorsal Attention Network | -5.99 | -2.01 | 0.106 | -0.88 | -0.26 | 0.922 |
| Ventral Attention Network | -9.12 | -2.1 | 0.106 | -15.36 | -3.41 | 0.020* |
| Limbic Network | 2.79 | 0.37 | 0.809 | 5.36 | 0.78 | 0.922 |
| Frontoparietal Network | -8.73 | -1.72 | 0.151 | 4.52 | 1.01 | 0.794 |
| Default Mode Network | 2.88 | 0.51 | 0.809 | 5.19 | 1.01 | 0.794 |

Note. The dependent variable in all models is the longitudinal change in self-control. B represents the unstandardized regression coefficient. Baseline Vulnerability examines whether intrinsic functional connectivity at Time 1 (T1 FC) predicts subsequent decline in self-control. Longitudinal Co-change examines whether longitudinal alterations in intra-network connectivities track concurrent self-control decline. To rigorously isolate true longitudinal covariation, all models were adjusted for baseline self-control, corresponding baseline functional connectivity, sex, and longitudinal head motion changes. The False Discovery Rate method was applied to correct for multiple comparisons (correction applied across the full 28 whole-brain connectome edges for the co-change models). \* *p* < 0.05.

In contrast, the longitudinal reconfiguration of intra-network integration exhibited highly specific co-change effects. To robustly test anatomical specificity, we expanded our target scan to encompass the entire macroscopic connectome (7 intra-network and 21 between-network edges). Employing rigorous change score models adjusted for baseline connectivity, baseline self-control, sex, and longitudinal head motion, the longitudinal increase in VAN intra-network connectivity emerged as the most robust predictor of self-control decline (B = −15.36, t = −3.41, *p*_FDR = 0.020; Table 2). Crucially, this was the only connection across the entire connectome to survive FDR correction. Between-network couplings did not yield significant longitudinal associations (all *p*_FDR > 0.49). This indicates that as the VAN’s internal connectivity strengthens over time, individuals experience a specific decline in self-control. Given the VAN’s role in bottom-up salience detection, this heightened stimulus-driven processing may continuously compete with the cognitive resources necessary for top-down self-regulation.

This highly specific longitudinal co-change is visually illustrated in Figure 3. The partial regression plot confirms that this robust negative association is a pervasive, sample-wide phenomenon rather than a statistical artifact driven by extreme outliers. This distributional evidence solidifies the Ventral Attention Network as the primary neural locus tracking the longitudinal erosion of cognitive control.

**Figure 3.**
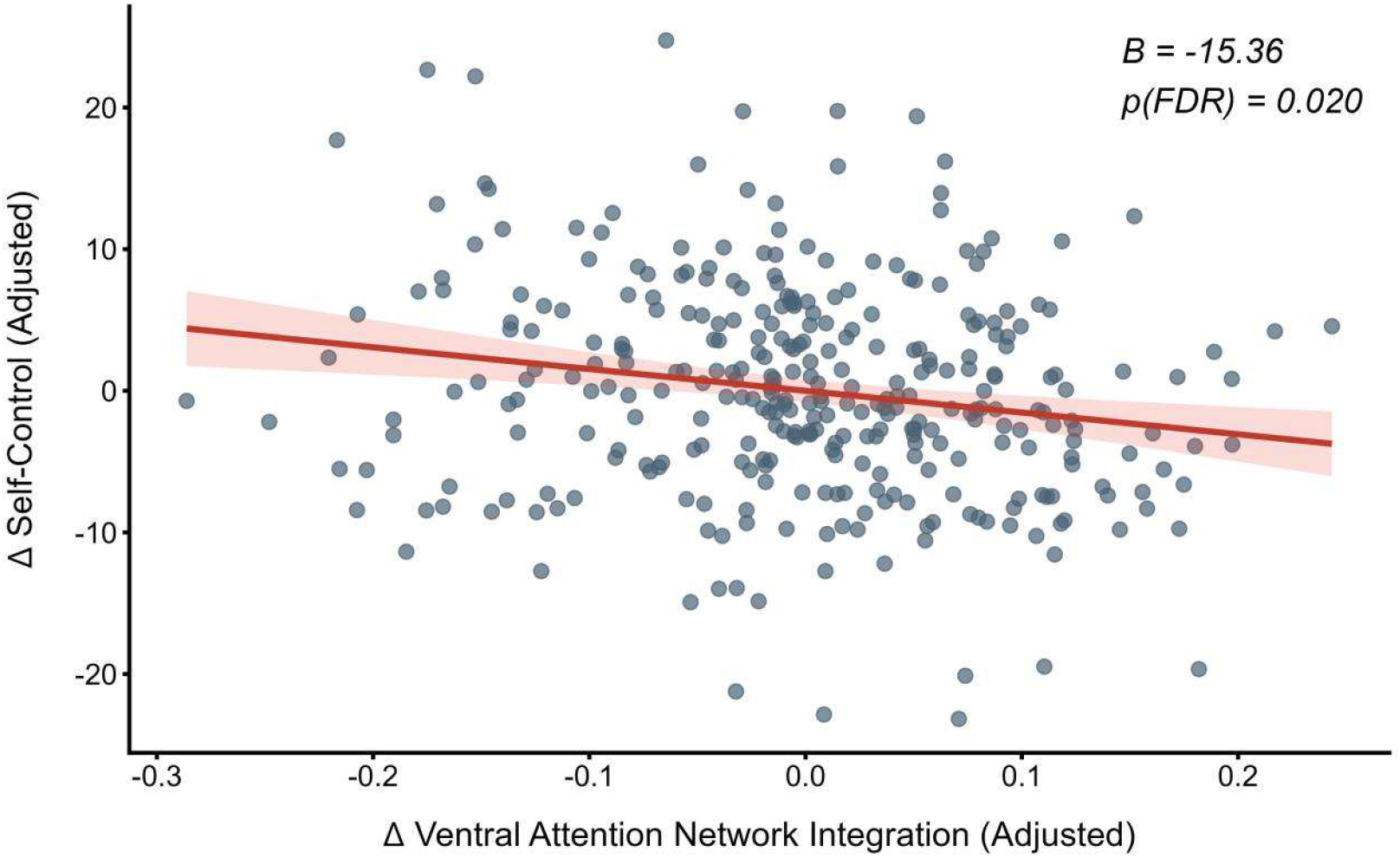
Longitudinal Increases in Ventral Attention Network Integration Negatively Predict Longitudinal Self-Control Trajectories. The scatter plot visualizes the robust negative association between longitudinal shifts in Ventral Attention Network integration (X-axis) and concurrent changes in self-control (Y-axis). The scatter plot displays the partial regression association, adjusting for baseline self-control, baseline network connectivity, sex, and longitudinal head motion changes. The solid red line depicts the unstandardized linear regression fit, and the shaded region represents the 95% confidence interval.

### 3.4. Longitudinal Decline in Self-Control Mediates the Link Between VAN Alterations and Heightened Sensitivities

To synthesize the longitudinal relationships among neural reconfiguration, cognitive resource decline, and behavioral symptom changes, we constructed a unified path analysis model. Specifically, we tested whether the longitudinal decline in self-control mediated the association between the longitudinal increases in intra-network connectivity of the Ventral Attention Network (ΔVAN) and the subsequent increases in reward and punishment sensitivities. To strictly isolate true longitudinal covariation, the model comprehensively adjusted for baseline self-control, corresponding baseline network connectivity, sex, and longitudinal head motion changes. The results are shown in Figure 4.

**Figure 4.**
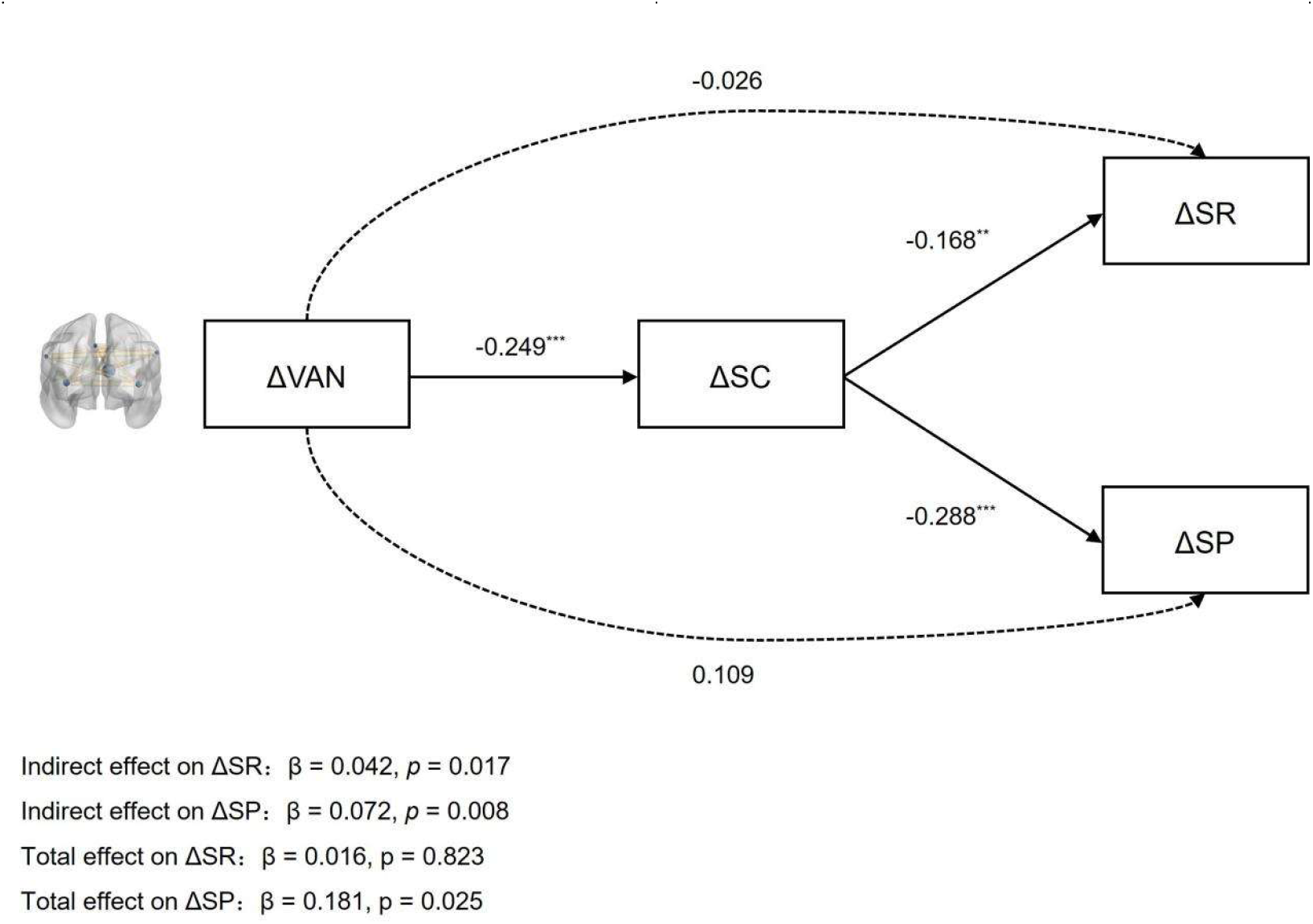
Longitudinal Path Model Linking Ventral Attention Network Alterations, Self-Control Decline, and Reward/Punishment Sensitivities. The path diagram illustrates the full mediation mechanism whereby the longitudinal increases in intra-network connectivity of the Ventral Attention Network predict changes in reward and punishment sensitivity via the longitudinal decline in self-control. Δ represents the raw longitudinal change scores from Time 1 to Time 2. Path values are standardized regression coefficients (β). Solid black lines denote significant pathways, while dashed grey lines represent non-significant direct effects. To rigorously isolate longitudinal dynamics, the model systematically adjusted for baseline self-control, baseline network connectivity, sex, and longitudinal head motion changes. ** *p* < 0.01, *** *p* < 0.001.

Consistent with our targeted univariate findings, the path analysis confirmed a highly significant behavioral pathway. The longitudinal increase in VAN integration significantly and negatively predicted the change in self-control (β = −0.249, *p* < 0.001), indicating that increased VAN intra-network connectivity strongly tracks a steeper decline in cognitive resources. In turn, the accelerated decline in self-control robustly predicted concurrent increases in both reward sensitivity (β = −0.168, *p* = 0.002) and punishment sensitivity (β = −0.288, *p* < 0.001). Crucially, the formal test of indirect effects revealed that increased VAN intra-network connectivity exerted a significant indirect impact on both behavioral phenotypes strictly through the decline in self-control. This indirect longitudinal pathway was significant for both reward sensitivity (Indirect effect: β = 0.042, *p* = 0.017) and punishment sensitivity (Indirect effect: β = 0.072, *p* = 0.008).

## 4. Discussion

This two-year longitudinal study integrated Latent Change Score Modeling (LCSM) with longitudinal resting-state neuroimaging to elucidate the temporal sequence underlying impulsivity, self-control, and reward/punishment sensitivities. Moving beyond traditional cross-sectional paradigms, our behavioral modeling first revealed a competitive framework: the longitudinal decline in self-control, rather than baseline or surging impulsivity, acts as the primary longitudinal predictor of the activation of trait sensitivities to both rewards and punishments. Building upon this behavioral foundation, our targeted neuroimaging analysis uncovered a unified full mediation model: the increased VAN intra-network connectivity is negatively associated with self-control capacity. Together, these findings establish a mechanistically grounded pathway that translates longitudinal intrinsic brain topological shifts into observable behavioral vulnerabilities in a healthy young adult cohort.

Behaviorally, the longitudinal competitive LCSM in the current study untangled the temporal interactions among the impulsivity-control-sensitivity triad. First, the model revealed that robust initial self-control exerted a significant protective effect against the longitudinal increases in impulsivity. This underscores self-control not merely as a concurrent state, but as a fundamental developmental buffer that restrains impulsive tendencies over time (Johnson, Voegtline, Ialongo, Hill, & Musci, 2023; Moffitt et al., 2011; June P Tangney, Boone, & Baumeister, 2018). Moreover, while traditional perspectives often frame trait impulsivity as the direct precursor to altered environmental sensitivities (Dawe & Loxton, 2004), the competitive longitudinal modeling in the current study revealed a distinct mechanistic hierarchy. When the longitudinal changes of impulsivity and self-control were pitted against each other to compete for variance, the longitudinal surge in reward and punishment sensitivities was neither predetermined by baseline impulsivity nor driven by concurrent impulsive spikes. Instead, the longitudinal decline in self-control emerged as the primary predictor of heightened sensitivities. This finding aligns with the framework of the strength model of self-control (Baumeister et al., 2007; Vohs et al., 2021).

Theoretically, self-control acts as the domain-general central executive “brake” modulating both the Behavioral Activation System (BAS, governing reward) and the Behavioral Inhibition System (BIS, governing punishment) (Corr, 2004; Hofmann, Schmeichel, & Baddeley, 2012). As these cognitive regulatory resources decline over time, the efficacy of this top-down buffer naturally diminishes. Consequently, individuals become more reactive to everyday environmental cues. This regulatory fatigue manifests as a dual susceptibility: an intensified pull toward immediate gratification (heightened BAS reactivity) alongside a sharpened sensitivity to aversive outcomes (heightened BIS reactivity) (Inzlicht & Schmeichel, 2012; Wagner, Altman, Boswell, Kelley, & Heatherton, 2013). This theoretical shift within the current longitudinal framework highlights that the longitudinal shifts in environmental sensitivities is driven less by an ascending impulsive trait, and fundamentally more by the reduction of psychological brakes. Crucially, this behavioral sequence necessitates an inquiry into the underlying macroscopic neural architecture: continuously correlate with this regulatory decline over time?

Notably, the targeted neuroimaging analysis successfully anchored this behavioral decline to a specific macroscopic neural configuration without invoking clinical pathology. While baseline static network topologies merely set a subtle vulnerability, the longitudinal reconfiguration—specifically the increased VAN intra-network connectivity—powerfully tracked the decline in self-control. Functionally anchored in the anterior insula and the inferior frontal gyrus, the VAN serves as the healthy brain’s primary bottom-up “circuit breaker,” designed to flexibly orient attention toward salient environmental cues (Corbetta & Shulman, 2002). Notably, within macroscopic cortical parcellations (Yeo et al., 2011), the VAN structurally encompasses and is functionally homologous to the Salience Network (SN) (Seeley et al., 2007). This anatomical convergence is theoretically paramount: it signifies that this system does not merely govern spatial attention, but actively computes the affective and motivational significance of incoming stimuli (Uddin, 2015). To interpret this negative coupling within a non-clinical cohort, we cautiously hypothesize that an over-reliance on bottom-up salient processing might compete with top-down regulatory networks. Theoretical frameworks suggest that the human brain operates with limited computational resources, wherein affective and cognitive systems often interact competitively (Pessoa, 2008). Viewed through this lens, a persistently heightened connectivity within the VAN/SN complex might relate to a redistribution of resources, potentially constraining the bandwidth available for top-down executive systems like the frontoparietal network (Menon, 2011). However, given that salience-related networks are also known to coordinate and recruit cognitive control processes, this resource-competition interpretation remains a speculative hypothesis that warrants further investigation using task-based designs. Consequently, rather than representing a structural deficit, the altered VAN connectivity may reflect a functional shift that relates to the reduction of the young adult’s self-control capacity over time.

The synthesis of these behavioral and longitudinal network alterations yielded a full mediation model (Figure 4), which delineates a robust temporal association among these variables. Notably, after accounting for the mediating role of self-control decline, the direct longitudinal effects of VAN alterations on behavioral sensitivities were rendered non-significant. This full mediation theoretically bridges a critical gap between network neuroscience and personality psychology (Markett, Montag, & Reuter, 2018): sensory-driven network configurations do not translate into complex trait alterations directly (Krakauer, Ghazanfar, Gomez-Marin, MacIver, & Poeppel, 2017). Rather, a persistently over-integrated bottom-up network disrupts behavioral homeostasis strictly by relating to changes in the central executive resource. This pathway provides robust empirical support for the premise that self-control is the indispensable functional bridge linking intrinsic macroscopic brain topology to how individuals experience and react to the external world (DeYoung et al., 2010; Friedman & Robbins, 2022).

Several limitations of the current study warrant consideration. First, the sample comprised predominantly university students, who are generally healthy and at a specific developmental stage of late adolescence to early adulthood. While this reduces the risk of confounding clinical pathologies that characterize older or unhealthy people, future research should replicate these findings across more diverse age groups and clinical cohorts to test the generalizability of the model. Second, our sample exhibited a marked sex imbalance (∼77% female), which may limit generalizability, and future studies should explore whether hormonal fluctuations influence resting-state functional connectivity measurements. Finally, this study relied on resting-state fMRI which limits causal inference. Future studies employing task-based fMRI and non-invasive brain stimulation targeting the VAN or executive networks could establish a more definitive causal link within this neurobehavioral pathway.

## 5. Conclusion

This two-year longitudinal study provides robust empirical evidence for a longitudinal neurobehavioral pathway connecting macroscopic brain network reconfiguration to cognitive development in emerging adulthood. Our behavioral modeling demonstrates that longitudinal increases in reward and punishment sensitivities are fundamentally driven by the progressive decline in self-control resources, rather than baseline or concurrent impulsivity. Neurally, confirmed by a comprehensive macroscopic connectome scan, this regulatory decline is specifically and powerfully anchored to longitudinal increases in intra-network connectivity of the Ventral Attention Network (VAN). We propose that during this critical developmental window, an over-integrated VAN may reflect a neurodevelopmental shift toward stimulus-driven processing that competes with top-down self-regulation. By validating this unified full mediation model, our study establishes increased VAN intra-network connectivity as a specific neural marker of cognitive erosion and highlights the preservation of self-control capacity as a critical mechanism for promoting adaptive neurocognitive resilience during the transition to adulthood.

## Data Availability

The de-identified longitudinal behavioral datasets, extracted resting-state functional connectivity matrices of the macroscopic brain networks, and all custom analytical scripts (in R) required to reproduce the findings of this study (including Latent Change Score Modeling, targeted network scans, and path analysis) have been publicly deposited in Zenodo (DOI: 10.5281/zenodo.24304008).

## Acknowledgments

This work was supported by research grants from the National Natural Science Foundation of China (31972906), Fundamental Research Funds for the Central Universities (SWU2209235), the Innovation Research 2035 Pilot Plan of Southwest University (SWUPilotPlan006), and the Open Research Fund of the State Key Laboratory of Cognitive Neuroscience and Learning (CNLZD2102).

## Authorship Contribution Statement

**Jingzhen He:** Conceptualization, Methodology, Formal analysis, Writing-original draft, Writing-review & editing. **Haichao Zhao:** Conceptualization, Methodology. **Xu Lei:** Conceptualization. **Jiang Qiu:** Methodology. **Tingyong Feng:** Conceptualization, Methodology. **Hong Chen:** Methodology. **Ofir Turel:** Writing-review & editing. **Qinghua He:** Conceptualization, Methodology, Data curation, Funding acquisition, Writing-review & editing.

## Notes

### Competing Interest Statement

The authors have declared no competing interest.

